# Distinct Regulation of History-dependent Responses by Two Cortical Interneuron Populations

**DOI:** 10.1101/129593

**Authors:** Elizabeth A.K. Phillips, Christoph E. Schreiner, Andrea R. Hasenstaub

**Affiliations:** Coleman Memorial Laboratory, University of California - San Francisco, San Francisco, CA 94158, USA; Neuroscience Graduate Program, University of California - San Francisco, San Francisco, CA 94158, USA; Department of Otolaryngology - Head and Neck Surgery, University of California - San Francisco, San Francisco, CA 94158, USA; Center for Integrative Neuroscience, University of California - San Francisco, San Francisco, CA 94158, USA; Kavli Institute for Fundamental Neuroscience, University of California - San Francisco, San Francisco, CA 94158, USA

**Keywords:** forward suppression, adaptation, spectrotemporal context, cortex, interneurons

## Abstract

Cortical responses to repeated stimuli are highly dynamic and rapidly adaptive. Such rapid changes are prominent in all sensory cortices, across which many aspects of circuitry are conserved. As an example, in the auditory cortex, preceding sounds can powerfully suppress responses to later, spectrally similar sounds – a phenomenon called forward suppression. Whether cortical inhibitory networks shape such suppression, or whether it is wholly regulated by common mechanisms such as synaptic depression or spike-frequency adaptation, is controversial. Here, we show that optogenetically suppressing somatostatin-positive interneurons reveals facilitation in neurons that are normally forward-suppressed. This is accompanied by a weakening of forward suppression, suggesting that these interneurons regulate the strength of forward interactions. In contrast, inactivating parvalbumin-positive interneurons does not change suppression strength, but does alter its frequency-dependence. These results establish a role of cortical inhibition in forward suppression and link specific aspects of rapid sensory adaptation to genetically distinct interneuron types.

## Introduction

Behaviorally meaningful sensory inputs, such as the sound of music or the wafting scent of a pie, rapidly change over time. In all sensory modalities, the dynamics of these sensory inputs influence both our perception of them and how the brain processes them. For instance, neural responses in the visual (Ohzawa et al., 1982), auditory (Ulanovsky et al., 2003), somatosensory (Simons, 1978), and olfactory (Wilson, 1998) cortices rapidly diminish to repeated presentations of the same stimulus. Such sensory adaptation is considered a mechanism for ignoring redundancies, detecting changes, and matching neural responses to the statistics of the sensory scene (Barlow, 1961; Fairhall et al., 2001); yet, the neuronal circuits that support these rapid response changes are not well-understood.

Throughout the auditory system, such adaptation is prominent and strongly depends on the history of recent stimulation. Neural responses to a sound are often completely abolished when preceded by a spectrally similar sound and more weakly suppressed if preceded by a spectrally dissimilar sound – a phenomenon called forward suppression (Brosch and Schreiner, 1997; Calford and Semple, 1995). This suppression is present as early as the auditory nerve (Harris and Dallos, 1979) and in multiple subcortical areas (Malone and Semple, 2001; Schreiner, 1981; Watanabe and Simada, 1971). However, responses in the auditory cortex tend to follow lower repetition rates (Creutzfeldt et al., 1980; Miller et al., 2002; Yao et al., 2015), and recover more slowly from sensory stimulation (Fitzpatrick et al., 1999), suggesting that forward suppression is, in part, cortically generated. Although it has been argued that forward suppression at the level of the inferior colliculus can account for some perceptual effects (Nelson et al., 2009), the temporal transformations within the cortex argue for further refinement of history-dependent processing.

How does such refinement occur at the cortical level? Diverse populations of specialized GABAergic interneurons interact to dynamically regulate neural activity. In the auditory cortex, their influence on spectral processing has been well established through direct intracellular recordings (Tan et al., 2004; Volkov and Galazjuk, 1991; Wehr and Zador, 2003), correlative studies in aged or presbycutic animals (Caspary et al., 2008), and application of GABA receptor agonists and antagonists (Kaur et al., 2004; Wang et al., 2002). However, their contribution to temporal processing is less evident. For example, in a recent study (Yao et al., 2015), application of GABA receptor antagonists did not attenuate forward suppression, suggesting that synaptic inhibition does not contribute. Moreover, other neural mechanisms, such as spike-frequency adaptation (Abolafia et al., 2011) and short-term synaptic depression (Bayazitov et al., 2013; Wehr and Zador, 2005), are pronounced in auditory cortex and in other sensory modalities, and have been hypothesized to dominate such history-dependent interactions (May et al., 2015). Assessing the role of inhibition is further complicated by the diversity of cortical interneurons, whose distinct contributions cannot be resolved by standard extracellular recording techniques or pharmacological methods and whose activity patterns are profoundly affected by both behavioral state (Fu et al., 2014; Steriade et al., 2001) and anesthesia (Adesnik et al., 2012; Haider et al., 2013). Thus, whether intracortical inhibition plays a role in forward suppression in awake animals, let alone how specialized inhibitory subpopulations might be involved, remains unresolved.

Here, in the auditory cortex of waking mice, we use optogenetics to test whether synaptic inhibition contributes to forward suppression and whether somatostatin-positive (Sst+) or parvalbumin-positive (Pvalb+) interneurons support history-dependent interactions in distinct ways.

## Results

### The history of auditory stimulation affects tone responses in diverse ways

We placed mice on a free-floating spherical ball, fixed their heads in place, and used 16-channel linear probes to record single units while the mice passively listened to sounds (Figure 1a). To measure the effects of stimulus history on auditory responses we presented 10-50 trials of a forward suppression stimulus: a 50 ms “masker” tone followed by a 50 ms “probe” tone separated by a 20 ms gap (stimulus-onset asynchrony: 70ms; Figure 1b). The masker tone varied in frequency across trials while the frequency of the probe tone remained fixed at the unit’s preferred frequency. Only units with a probe frequency within 0.5 octaves of the preferred frequency were analyzed. To measure the baseline response to the probe tone (without masking), on randomly interleaved trials (probe alone (PA) trials), the masker tone was omitted. This allowed us to assess the response to the probe tone alone, and how preceding tones of varying frequency altered its response.

**Figure 1:**
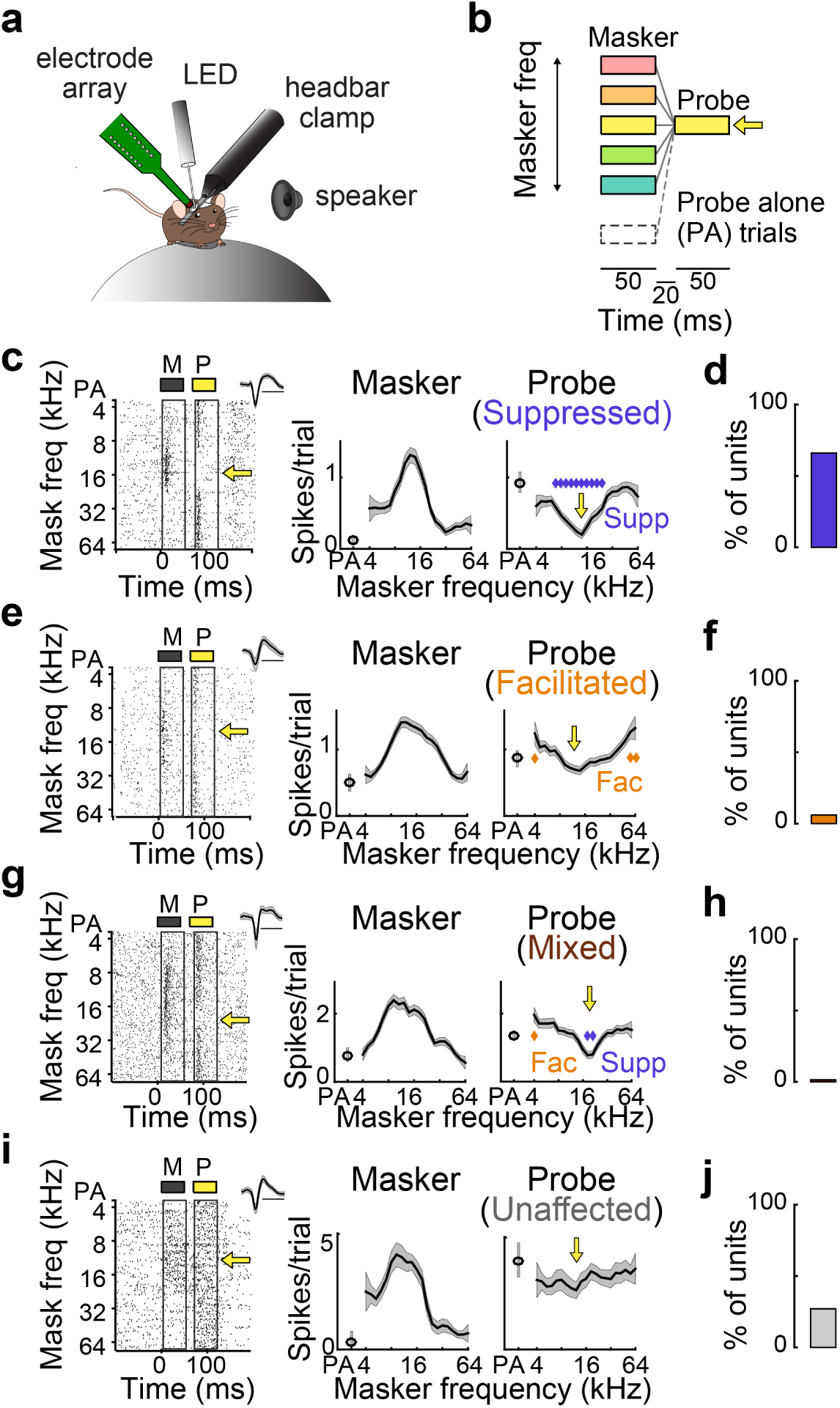
Prior tones affect responses to later tones in diverse ways. **a)** Recordings are performed from the auditory cortex of awake mice, headfixed atop an air-floated spherical treadmill. **b)** Forward suppression stimulus: a masker tone (varied frequency) precedes a probe tone (fixed frequency, indicated by yellow arrow). On probe alone (PA) trials the masker tone is omitted. **c)** Left: single unit response raster to the masker (gray rectangle) and probe (yellow rectangle), as a function of masker frequency. Inset shows spike waveform (scale bar: 1 ms). Right: responses (mean ± SEMs) to the masker and probe, as a function of masker frequency. Blue diamonds: significantly suppressed probe responses. Yellow arrow: probe frequency. **d)** 64% of units were suppressed. **a) e)** As (**c**) for a forward-facilitated unit. Gold diamonds: facilitated probe responses. **f)** 13% of units were facilitated. **g)** As (**c**) for a unit with a mixed effect. Blue: diamonds: suppressed responses; gold diamond: facilitated response. **h)** 3% of units had mixed effects. **i)** As (**c**) for an unaffected unit. **j)** 20% of units were unaffected.

The presence of a masker tone had multiple, qualitatively different effects on the response to the probe tone. The majority of units (43/67, N = 20 mice) exhibited forward suppression (Figure 1c,d). A minority of units (9/67) showed forward facilitation, which generally occurred when the masker and probe frequencies were dissimilar (Figure 1e,f). An even smaller portion of units (2/67) showed mixed effects, i.e. both forward suppression and forward facilitation (Figure 1g,h), and the remainder (13/67) were unaffected by the presence of a preceding masker stimulus (Figure 1i,j).

### Altering inhibition changes the quality of forward interactions

To directly test whether synaptic inhibition within the cortex contributes to forward suppression, and whether Sst+ and Pvalb+ interneuron populations differentially support such adaptation, we used two mouse lines in which we could optogenetically inactivate either Sst+ or Pvalb+ interneurons (Ai35/Sst-Cre and Ai35/PV-Cre crosses, respectively; see Methods). On randomly interleaved presentations of the stimulus, we inactivated either of the two interneurons by shining green light on the surface of the auditory cortex while recording from single units in the auditory cortex. To avoid direct effects of the optogenetic inactivation in our analysis we excluded units for which green light significantly suppressed masker-evoked firing rates (Ai35/Sst-Cre: 2/30; Ai35/PV-Cre: 4/37).

We found that inactivating Sst+ interneurons in Ai35/Sst-Cre mice (N = 11) considerably altered the way that prior stimuli affected responses to the probe tone (Figure 2; Supplemental figure 2). Of the units that were forward-suppressed, two thirds became either facilitated (4/18), mixed (2/18), or unaffected (6/18) with inactivation of Sst+ cells, while the other third (6/18) remained purely suppressed. Neurons unaffected by a masker in the light-off condition became either suppressed (3/7), facilitated (2/7), or remained unaffected (2/7). In total, the result of inactivating Sst+ interneurons was a net decrease in the number of suppressed units and a net increase in the number of facilitated, mixed, and unaffected units (Figure 2g). In contrast, inactivating Pvalb+ interneurons in Ai35/Pvalb-Cre mice (N = 9) did not alter the quality of forward interactions in most forward-suppressed units; only one unit became unaffected while the majority (20/21) remained purely suppressed. Interestingly, the majority of unaffected units (7/10) became forward-suppressed with inactivation of Pvalb+ cells. Overall, inactivating Pvalb+ cells resulted in a net increase in the number of suppressed units and a net decrease in the number of facilitated and unaffected units (Figure 2n). These distinct effects of Sst+ and Pvalb+ interneuron inactivation were not explained by differences in overall firing rate changes, as inactivation of Sst+ and Pvalb+ interneurons produced similar changes in spontaneous (ranksum, p = 0.88), masker-evoked (ranksum, p = 0.63), and probe-evoked (ranksum, p = 0.23) firing rates (Supplemental Figure 1). These results demonstrate that intracortical inhibition affects the way in which stimulus history influences neural responses, and in particular, inhibition from Sst+ and Pvalb+ interneurons contributes to history-dependence in qualitatively different ways.

**Figure 2:**
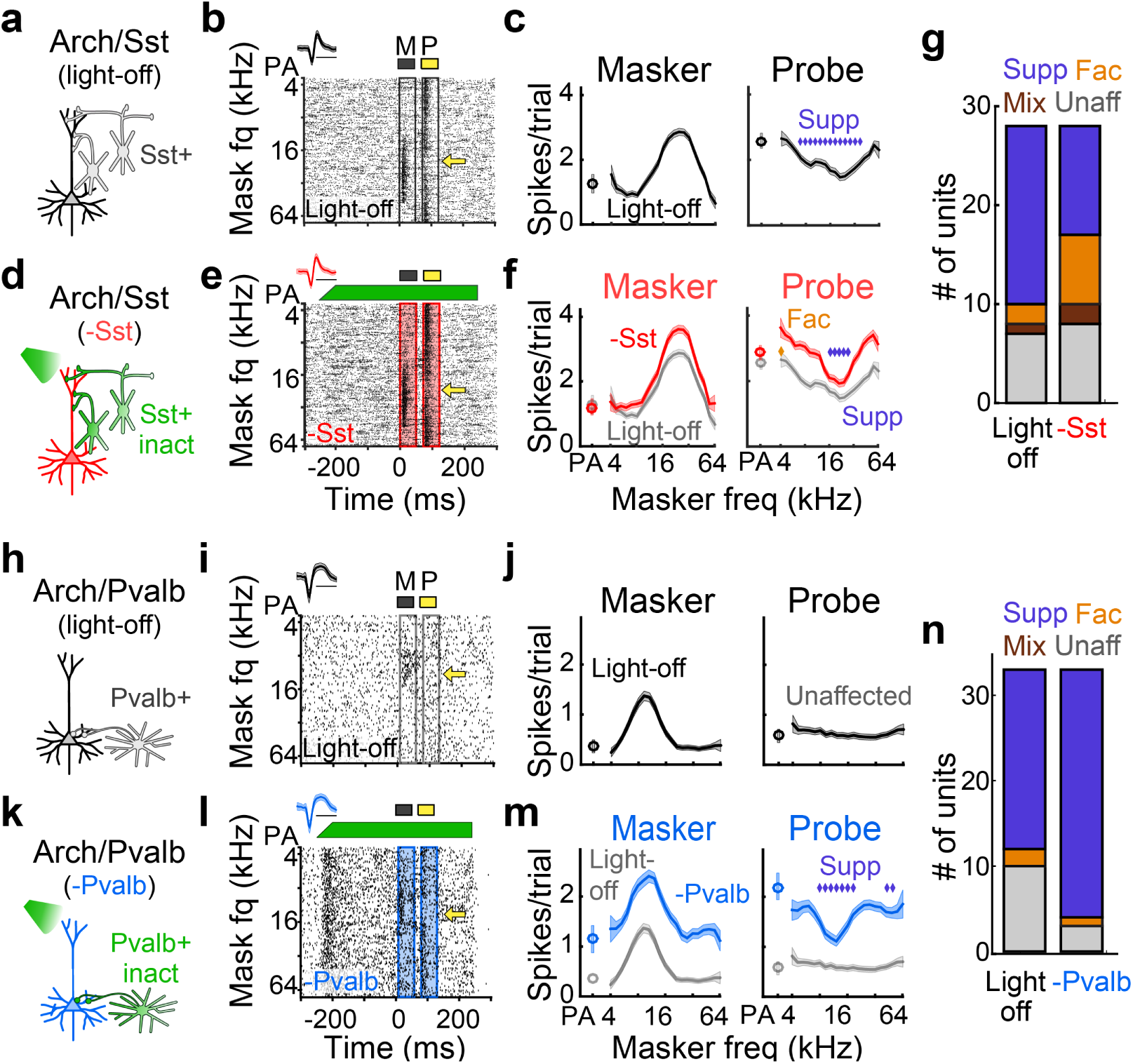
Inactivation of Sst+ versus Pvalb+ interneurons differentially alters the quality of forward interactions. **a)** Schematic of circuit during light-off trials. **b)** Single unit response raster from light-off trials. Yellow arrow: probe frequency. Inset shows spike waveform (scale bar: 1ms). **c)** Responses (mean ± SEMs) to the masker and probe, as a function of masker frequency, in light-off trials. Blue diamonds: significantly suppressed probe responses. **d)** Schematic of circuit during light-on trials, in which Sst+ cells are inactivated with green light. **e)** As (**b**) for light-on trials. Green bar: duration and power of the light. **f)** As (**c**) for Sst+ inactivation trials (red) versus light-off trials (gray). Sst+ inactivation produces facilitation of the probe response. Blue diamonds: suppressed probe responses; gold diamonds: facilitated probe responses. **g)** Proportion of suppressed, facilitated, mixed, and unaffected units without (left) and with (right) inactivation of Sst+ cells. **h-m)** As (**a-f**) for a single unit in which inactivation of Pvalb+ cells (blue) reveals forward suppression. **n)** As (**g**) for inactivation of Pvalb+ cells.

### Altering inhibition changes the relative strength and spectral dependence of forward suppression

In addition to these qualitative changes in forward interactions, does decreasing inhibition from Sst+ or Pvalb+ interneurons change the relative strength or shape of forward suppression? For units that exhibited forward suppression (including both suppressed and mixed units), we examined the relative strength of forward suppression by dividing the suppression profiles by the probe alone responses (Figure 3a). We performed this normalization separately for light-on and light-off trials to compare the strength of forward suppression with and without inactivation of either interneuron type. Among all forward-suppressed units in Ai35/Sst-Cre mice (n = 19 units from 9 mice), we found that inactivating Sst+ interneurons produced normalized suppression profiles that were shifted upwards compared to the normalized suppression profiles in light-off trials, i.e. inactivating Sst+ interneurons attenuated the strength of forward suppression approximately equally across all masker frequencies (Figure 3b). In contrast, among all forward-suppressed units in Ai35/Pvalb-Cre mice (n = 21 units from 8 mice), normalized suppression profiles with inactivation of Pvalb+ interneurons were generally similar to those from light-off trials (Figure 3c), suggesting that inactivating Pvalb+ cells did not attenuate the strength of forward suppression.

**Figure 3:**
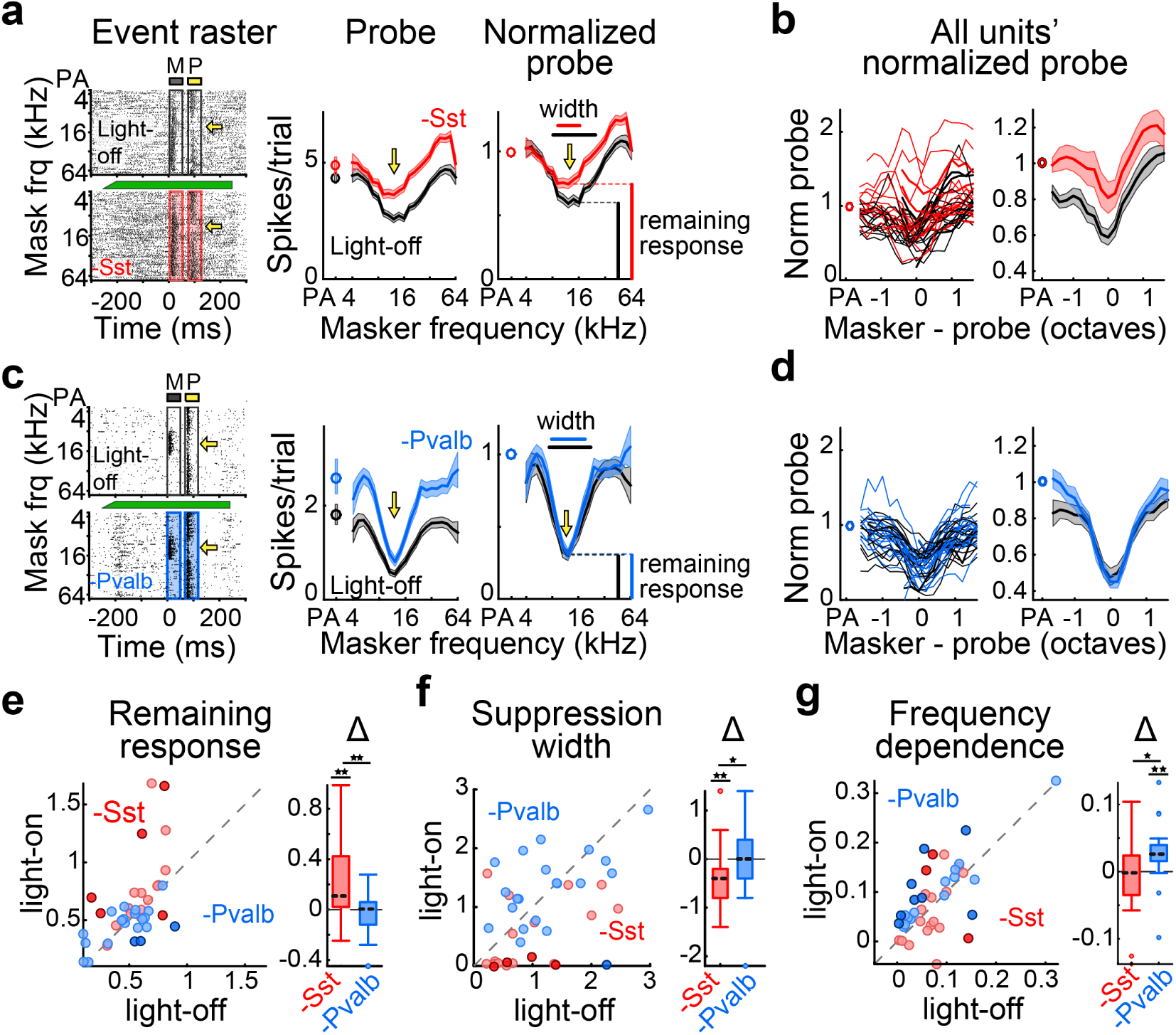
Inactivation of Sst+ versus Pvalb+ interneurons differentially alters the strength and shape of forward suppression. **a)** Left: Example single unit responses to masker and probe stimuli without (top) and with (bottom) inactivation of Sst+ cells. Middle: Responses (mean ± SEMs) to the probe as a function of masker frequency without (black) and with (red) inactivation of Sst+ cells. Right: Normalized probe responses without (black) and with (red) inactivation of Sst+ cells. Remaining response at probe frequency and suppression width are shown. **b)** All single units’ normalized probe responses (left) and unit-averaged probe responses (right), as a function of masker – probe distance, without (black) and with (red) inactivation of Sst+ cells. **c)** As (**a**) for an example single unit with inactivation of Pvalb+ interneurons (blue). **d)** As (**b**) for all units with inactivation of Pvalb+ interneurons. **e-g)** Remaining response at probe frequency (**e**), suppression width (**f**), and frequency dependence (**g**) without versus with inactivation of Sst+ (red) or Pvalb+ (blue) interneurons. Darker circles represent units with significant effects. Box and whisker plots show distribution of effect sizes (*p<0.05, **p<0.01, ***p<0.001).

To quantify these effects, we measured three aspects of units’ suppressed probe responses (Figure 3a,c). Firstly, the remaining response at probe frequency was defined as the normalized probe response when the probe was preceded by a masker of the same frequency (often the most suppressed probe response). Secondly, suppression width was measured as the range (in octaves) of masker frequencies that significantly suppressed probe responses below the probe alone response. Lastly, the frequency dependence of suppression was defined as the proportion of variance in the probe response that was explained by the frequency of the masker (ω^2^; see Methods).

Inactivation of Sst+ cells significantly increased remaining responses at probe frequency (signrank, p = 0.0048; Figure 3e) and decreased suppression widths (signrank, p =0.0093; Figure 3f), but did not change the proportion of variance explained by masker frequency (signrank, p = 0.69; Figure 3g), consistent with the idea that Sst+ inactivation weakens forward suppression evenly across all masker frequencies. On the other hand, inactivation of Pvalb+ interneurons did not affect remaining responses at probe frequency (signrank, p = 0.66; Figure 3e) or suppression widths (signrank, p = 0.66; Figure 3f), but did significantly increase the proportion of variance explained by the masker frequency (signrank, p = 0.0041; Figure 3g), consistent with the notion that Pvalb+ inactivation alters the frequency dependence of probe suppression. These results imply that Sst+ interneurons regulate the strength, but not the spectral dependence, of forward suppression. Pvalb+ interneurons, on the other hand, appear to be more important for regulating the spectral dependence, but not the overall strength, of forward suppression.

### Changes in tuning profiles do not account for changes in suppression profiles

One potential explanation for these observed changes in the strength of forward suppression is that inactivating interneurons indirectly alters forward suppression strength by increasing spiking activity in response to the masker, which could increase intrinsic adaptation currents, such as voltage- or potassium-gated potassium currents (Brown and Adams, 1980; Madison and Nicoll, 1984). To gain intuition for whether changes in the probe response were caused by increases in masker spiking activity, we selected trials from the light-off and light-on conditions with the identical numbers of spikes in response to the masker (Figure 4a-c). After selecting these masker-matched trials, the normalized suppression profiles with inactivation of Sst+ or Pvalb+ cells appeared highly similar to those across the whole set of trials (Figure 4d-e). Moreover, inactivation of Sst+ cells still increased the remaining responses at probe frequency (signrank, p = 0.0022), while inactivation of Pvalb+ cells still increased the frequency dependence of suppression (signrank, p = 0.0015). Sst+ inactivation no longer decreased suppression widths (signrank, p = 0.58), however, most likely because the decrease in number of trials reduced the likelihood that responses would be significantly suppressed. Overall, these results suggest that the effects of inactivating Sst+ or Pvalb+ interneurons on forward suppression are not explained by changes in the firing rate of the neuron preceding the presentation of the probe tone.

**Figure 4:**
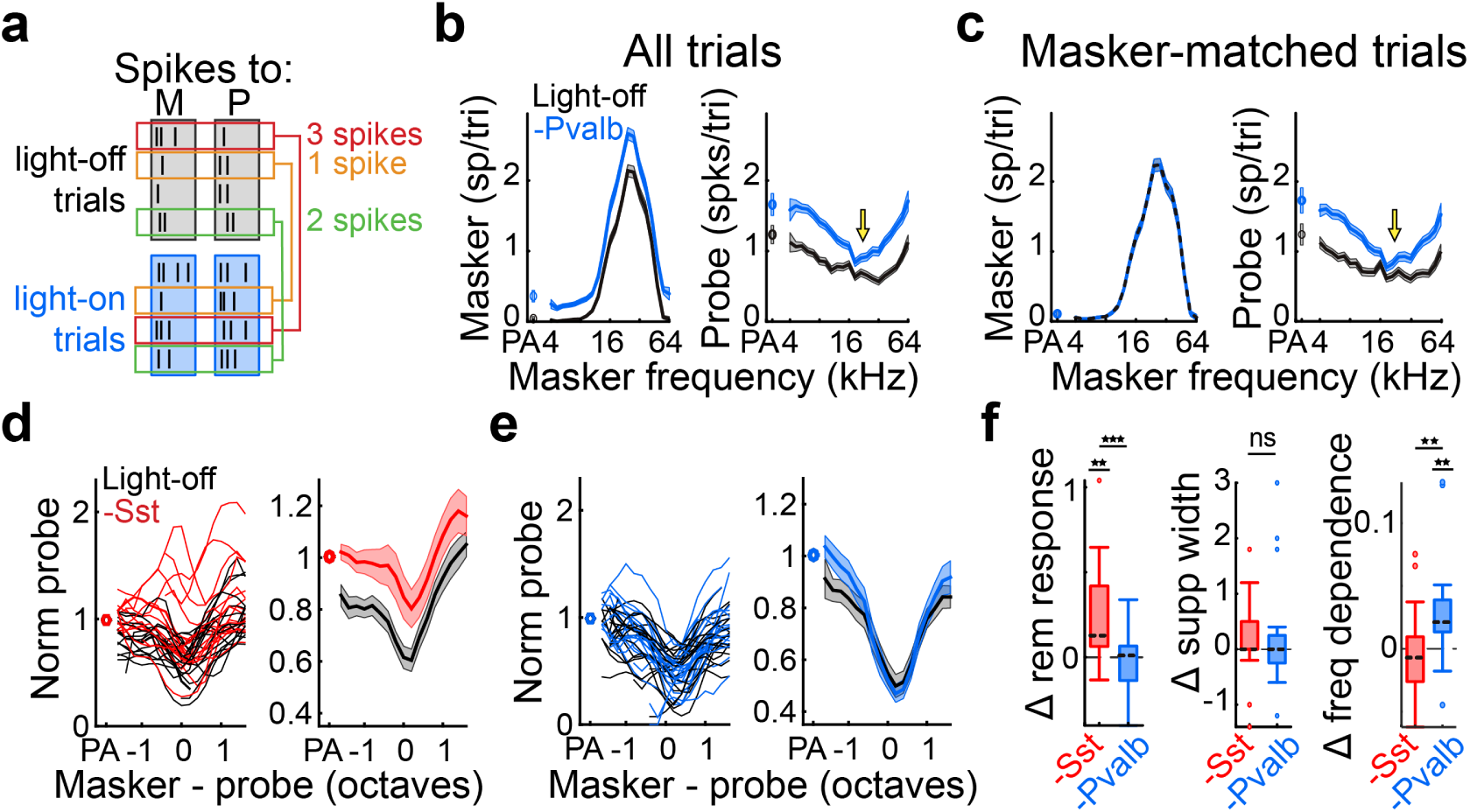
Effects of Sst+ and Pvalb+ inactivation on suppression are not explained by changes in masker responses. **a)** We select trials from the light-off and light-on conditions that contain identical numbers of masker-evoked spikes. **b)** Example unit responses (including all trials) to the masker (left) and probe (right) with (blue) and without (black) inactivation of Pvalb+ interneurons. **c)** As (**b**) for trials that are matched based on masker responses. **d)** All units’ (left) and unit-averaged (right) normalized probe responses from masker-matched trials, as a function of masker – probe distance, without (black) and with (red) inactivation of Sst+ cells. **e)** As (**d**) with inactivation of Pvalb+ interneurons. **f)** Changes in remaining response, suppression width, and frequency dependence with inactivation of Sst+ (red) or Pvalb+ (blue) interneurons (*p<0.05, **p<0.01, ***p<0.001).

## Discussion

In the auditory cortex of waking mice, we find that the influences of earlier sounds on responses to subsequent sounds (called forward suppression) are shaped by cortical synaptic inhibition, and that Sst+ and Pvalb+ interneurons support separate aspects of this adaptive process. Optogenetic inactivation of Sst+ interneurons during tone sequences weakened forward suppression, whereas inactivating Pvalb+ interneurons altered its spectral dependence. Additionally, these interneuron-specific effects on forward suppression were not caused by a change in the response to the earlier tone, as matching first-tone responses between light-off and light-on trials did not alter the effects on forward suppression. Overall, we conclude that cortical inhibition does regulate forward suppression, and that Sst+ interneurons control the strength of forward suppression, potentially influencing the detectability of later stimuli, whereas Pvalb+ interneurons regulate its dependence on the frequency of prior stimulation, potentially altering the discriminability of tone sequences.

This role for cortical inhibition in forward suppression has not been observed before. For example, using longer stimulus-onset asynchronies, Wehr and Zador (2005) demonstrated in anesthetized rats that inhibitory conductances in auditory cortex do not last longer than 100 ms, and therefore cannot explain the long-lasting neural and behavioral effects of forward suppression. Thus, whether synaptic inhibition contributes to forward suppression may depend on the timing between stimuli. However, ketamine anesthesia used in Wehr and Zador (2005) may also explain some of this effect by potentially decreasing the recruitment of synaptic inhibition in ways that may differ between interneuron types (Quirk et al., 2009). More recently, Yao et al. (2015) pharmacologically infused GABAa or GABAb receptor antagonists into the auditory cortex and found no attenuation of forward suppression. However, GABA receptor antagonists act upon all sources of synaptic inhibition, potentially conflating the distinct effects of different interneurons. On the other hand, optogenetic manipulation of interneurons has its own set of caveats, including counterintuitive network effects (Phillips and Hasenstaub, 2016; Seybold et al., 2015) and complex interactions among connected interneuron networks (Pfeffer et al., 2013).

What are the potential mechanisms by which inactivation of Sst+ or Pvalb+ interneurons produce different effects on forward suppression? One possibility is that manipulating inhibition has an indirect effect on forward suppression by acting through activity-dependent mechanisms, such as synaptic depression or spike-frequency adaptation. For example, suppressing inhibition increases local network activity, increasing synaptic or spike-frequency adaptation within the network and leading to significant changes in both sensory responsiveness (Hasenstaub et al., 2007) and the strength of forward suppression (Castro-Alamancos, 2004). These effects, however, are not large contributors to forward suppression because Sst+ and Pvalb+ inactivation similarly increased local network activity (Supplemental figure 1) yet differentially altered forward suppression.

Another potential mechanism is that Sst+ and Pvalb+ interneurons are recruited by feedforward inputs with different dynamics. In both somatosensory and auditory cortical slices, thalamocortical inputs onto Sst+ interneurons facilitate when stimulated at rates similar to the onset asynchrony of the tones presented in the current study [10-60 Hz; (Beierlein et al., 2003; Takesian et al., 2013; Tan et al., 2008)]. Thus, Sst+ cells may be robustly and persistently activated during repeated sensory stimulation, allowing them to directly suppress their targets across multiple stimulations. Pvalb+ interneurons, on the other hand, receive depressing thalamocortical inputs in slice and are only activated transiently (Beierlein et al., 2003; Takesian et al., 2013; Tan et al., 2008), diminishing their ability to suppress their targets across repeated stimulations. Consequently, Pvalb+ interneurons may only suppress responses to successive stimuli when not strongly activated by the first stimulus (for instance, if the stimulus lies beyond the edges of their receptive fields). This could explain why inactivating Pvalb+ cells produced a minor attenuation of forward suppression at the edges of the suppression curve, but not at the center (Figure 3d), and increased the frequency dependence of suppression. These known differences in synaptic dynamics between Sst+ and Pvalb+ interneurons, in combination with the current result that inactivation of Sst+ versus Pvalb+ interneurons differentially alters forward suppression, offer a potential mechanism by which inhibition from different inhibitory microcircuits support distinct, non-overlapping aspects of temporal processing within the cortex.

Context-dependent effects occur in other sensory modalities as well, such as contrast adaptation in visual cortex and adaptation of whisker deflection responses in somatosensory cortex. Although studies suggest that synaptic depression and spike-frequency adaptation dominate these effects, much of this work was either conducted in quiet slices (Castro-Alamancos and Connors, 1997; Sanchez-Vives et al., 2000; Varela et al., 1997), where synaptic depression is pronounced, or under anesthesia (Chung et al., 2002; Freeman et al., 2002; Nelson, 1991), in which interneurons’ effects are distorted. By performing selective manipulation experiments in awake animals, we demonstrate a previously unidentified contribution of synaptic inhibition to forward suppression in the auditory cortex, perhaps translating to context-dependent interactions in other cortical areas, which the field of systems neuroscience is already beginning to discover (Adesnik et al., 2012; Fu et al., 2014; Natan et al., 2015).

## Methods

### Protocols

All experiments were performed in accordance with the National Institutes of Health guidelines and were approved by the IACUC at the University of California, San Francisco.

### Animals

Adult mice (male or female) were housed under a 12 hr/12 hr light/dark cycle and used for experiments between the ages of 6 and 12 weeks old. To target either Sst+ or Pvalb+ cells for optogenetic manipulation, we crossed Sst-IRES-Cre or Pvalb-IRES-Cre knock-in lines (JAX stock no. 013044 and 008069, respectively) to the Ai35 line (JAX stock no. 012735), which encodes the light-gated proton pump Archaerhodopsin-3 (Arch) fused to GFP under the CAG promoter after a loxP-flanked STOP cassette.

### Surgeries

Animals were induced under anesthesia with 3% isoflurane, maintained with 1.25-2% isoflurane, and injected with pre- and post-operative multimodal analgesics. A custom metal headbar was affixed over the right temporal skull with dental cement, and a silicone elastomer was placed over the exposed bone. Animals recovered for 2-5 days after which a craniotomy was drilled above the right auditory cortex, and new silicone elastomer was placed over the exposed brain. Animals recovered for 1-3 hours before recordings.

### Data acquisition and stimuli

Animals were head-fixed above an air-floated spherical treadmill (Niell and Stryker, 2010) and the silicone plug was removed. A 16 site linear probe (50 µm spacing, Neuronexus) was inserted approximately perpendicular to the cortical surface. Neural activity was amplified, digitized, and recorded continuously at 24,414 Hz with Tucker-Davis Technologies hardware.

Auditory stimuli were generated in MATLAB (MathWorks) and presented through a free-field speaker (ES1, Tucker-Davis Technologies) directed toward the mouse’s left ear. To determine the best frequency of the units in the recording site, 50 ms tones (4 kHz to 64 kHz, 0.2 octave spacing; 0-60 dB, 5 dB increments) were presented. The best frequency of the recording site at 10-15 dB above threshold was used as the frequency of the probe tone for subsequent forward suppression experiments. The forward suppression stimulus consisted of two sequential tones, a 50 ms masker tone followed by a 50 ms probe tone separated by a 20 ms gap (stimulus-onset asynchrony of 70ms). The probe remained constant at the best frequency, 10-15 dB above threshold, while the masker tone randomly varied in frequency (4 kHz to 64 kHz, 0.2 octave spacing) at 15-20 dB above threshold. On a random subset of trials, called probe alone (PA) trials, the probe was presented without a preceding masker. Only units for which the probe frequency was within 0.5 octaves of the best frequency were analyzed.

On randomly interleaved trials green light was shined directly above the surface of auditory cortex through a 400 micron fiber. The light turned on 250 ms before sound onset, and the power linearly ramped upwards for 50 ms before reaching maximum power (10-15 mW). After sound offset, the light remained on for an additional 120 ms, totaling 490 ms.

### Constructing single unit tuning and suppression profiles

Neural events that crossed a threshold of 4 standard deviations above background were sorted into single units using custom software (MATLAB code by Matthew Fellows, based on (Calabrese and Paninski, 2011)). For each single unit (n = 81) we constructed PSTHs by collapsing events across masker frequency and grouping events into 1 ms bins, separately for light-off and light-on conditions. Units were defined as auditory responsive if the activity during the masker tone (50 ms) was significantly (α = 0.01, rank-sum test) greater than in the 50 ms before masker onset, in the light-off condition. For all auditory responsive units (n = 77), we defined the masker response latency as the time at which masker-evoked activity crossed 3SDs above spontaneous activity. Frequency tuning profiles were calculated as the average firing rate as a function of masker frequency during the 50 ms period after masker response latency. Units were defined as tuned if the tuning profile was significantly (α = 0.05, Kruskal-Wallis) modulated by frequency in the light-off condition.

For all tuned units (n = 67), we then generated suppression profiles, separately for light-on and light-off conditions, by measuring the average firing rate as a function of masker frequency during the 50 ms period after probe response onset (taken to be the same as masker response onset). Normalized suppression profiles were generated by dividing light-on and light-off suppression profiles by the PA response in the light-on and light-off conditions, respectively.

### Forward suppression classification and quantification

Classifications of suppression curves were performed separately for the light-off and light-on conditions. Units were suppressed if at least one of the masker stimuli significantly (α = 0.01, rank-sum) suppressed the probe response relative to the PA response. Units were facilitated if at least one masker stimulus significantly (α = 0.01, rank-sum) increased the probe response relative to the probe alone response. Units were mixed if they exhibited both suppression and facilitation, and units were unaffected if masker stimuli had no effect on the response to the probe. Remaining response at probe frequency was measured as the normalized firing rate of the probe response when the masker and probe were the same frequency. Suppression width was quantified as the range of masker frequencies, in octaves, that suppressed the probe response relative to the PA response. Frequency dependence was measured as *ω*^2^, an unbiased measure of the proportion of variance in the probe response that was explained by the frequency of the masker. To calculate *ω*^2^, we performed a one-way ANOVA between the probe response and masker frequency and computed the sum of squares between masker frequencies (*SS*_*fffq*_), the total sum of squares (*SS*_*total*_), the mean square error (*MS*_*error*_), and the degrees of freedom (*df*). *ω*^2^ was then defined as:

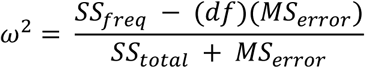

